# Proxi-RIMS-seq2 applied to native microbiomes uncovers hundreds of known and novel ^m5^C methyltransferase specificities

**DOI:** 10.1101/2024.07.15.603628

**Authors:** Weiwei Yang, Yvette Luyten, Emily Reister, Hayley Mangelson, Zach Sisson, Benjamin Auch, Ivan Liachko, Richard J. Roberts, Laurence Ettwiller

## Abstract

Methylation patterns in bacteria can be used to study Restriction-Modification (RM) or other defense systems with novel properties. While ^m4^C and ^m6^A methylation is well characterized mainly through PacBio sequencing, the landscape of ^m5^C methylation is under-characterized. To bridge this gap, we performed RIMS-seq2 on microbiomes composed of resolved assemblies of distinct genomes through proximity ligation. This high-throughput approach enables the identification of ^m5^C methylated motifs and links them to cognate methyltransferases directly on native microbiomes without the need to isolate bacterial strains. Methylation patterns can also be identified on viral DNA and compared to host DNA, strengthening evidence for virus-host interaction. Applied to three different microbiomes, the method unveils over 1900 motifs that were deposited in REBASE. The motifs include a novel 8-base recognition site (CAT^m5^CGATG) that was experimentally validated by characterizing its cognate methyltransferase. Our findings suggest that microbiomes harbor arrays of untapped ^m5^C methyltransferase specificities, providing insights to bacterial biology and biotechnological applications.

## Introduction

The role of methylation, an important epigenetic mark, extends beyond the well-known Restriction-Modification (RM) systems in bacteria, playing roles in the orchestration of gene expression and other cellular mechanisms (Anton and Roberts 2021). While ^m4^C and ^m6^A methylation patterns have been extensively identified, particularly through the use of Pacific Biosciences single molecule sequencing, our understanding of the landscape of ^m5^C methylation and methyltransferase specificities in bacteria lags behind. Efforts utilizing Tet-assisted PacBio sequencing (Clark et al. 2013) and more recently Nanopore sequencing technology have begun to bridge this gap (Tourancheau et al. 2021). However, these methods require substantial amounts of native DNA and have limited throughput, posing significant hurdles.

Conversely, high-throughput short-read sequencing enables the rapid sequencing of entire microbiomes at very high coverage, expanding the study of microbiomes to a broad range of starting materials and versatile selection protocols. In addition to sequencing genomes, methylome information can be obtained through short-read sequencing. However, the most common approach to overlay methylation to sequence information involves deamination of cytosine to uracil using bisulfite or enzymatic treatments. These treatments significantly alter the DNA sequence, making them incompatible with mixtures of non-model organisms typically observed in microbiomes for which no reference genomic sequences are available.

Recently, RIMS-seq (Baum et al. 2021) has been developed to sequence non-model organisms, such as bacterial genomes, and simultaneously determine ^m5^C methyltransferase specificities without requiring a reference genome. However, RIMS-seq is limited in its application to native microbiomes because short-read assemblies do not result in long enough contigs to assemble complete genomes from complex metagenomes.

To address this complexity, proximity ligation metagenomic sequencing has emerged as a strategy to deconvolute complex microbial communities ((Marbouty et al. 2014), (Burton et al. 2014), (Bickhart et al. 2022). This technique involves crosslinking DNA within intact cells prior to lysis, preserving the spatial information about sequences originating within the same cells. This information is recovered through ligating digested ends centered on a crosslink site, and the resulting chimeric molecules are analyzed via paired-end sequencing. When combined with a metagenomic assembly, these chimeric reads can be used to link contigs from the same consensus genome together into very high quality bins (referred throughout the manuscript as metagenome-assembled genomes or MAGs), and additionally to associate mobile elements like phages and plasmids with their microbial hosts without culturing.

In response to these challenges, we developed Proxi-RIMS-seq2 which combines an improved version of RIMS-seq and proximity ligation technologies to simultaneously assess the genetic and ^m5^C epigenetic information on genome-resolved microbiomes. To exemplify the potential of this methodology, we applied Proxi-RIMS-seq2 to three highly diverse microbiomes and linked the methylated motifs to their cognate methyltransferases.

## Result

### RIMS-seq2 accurately identifies methylated context on a mock gut microbiome

Earlier versions of RIMS-seq and RIMS-seq2 were employed on mock microbiomes and human genomic DNA, respectively (Baum et al. 2021; Yan, Wang, and Ettwiller 2024). We first tested RIMS-seq2 on the mock gut microbiome (ATCC ® MSA-1006™) spiked with XP12 bacteriophage genomic DNA, where all cytosines have been replaced by ^m5^C. RIMS-seq2 has a higher deamination rate compared to RIMS-seq and would require significantly less coverage per genome. Sequencing reads were downsampled to 1 Million Paired-end reads and were mapped to the mock community reference genomes. We calculated the imbalance of C-to-T transition between paired-end read 1 and 2, established previously to be linearly correlated with methylation ((Baum et al. 2021; Yan, Wang, and Ettwiller 2024). Using XP12 as our deamination control, we achieved around 1.42% deamination rate on ^m5^C which is consistent with the previous report of RIMS-seq2 (Yan, Wang, and Ettwiller 2024) (**Supplementary Figure 1A**). This deamination level is large enough to detect the m5C methylase specificity but too low to affect sequencing and assembly quality.

**Supplementary Figure 1:**
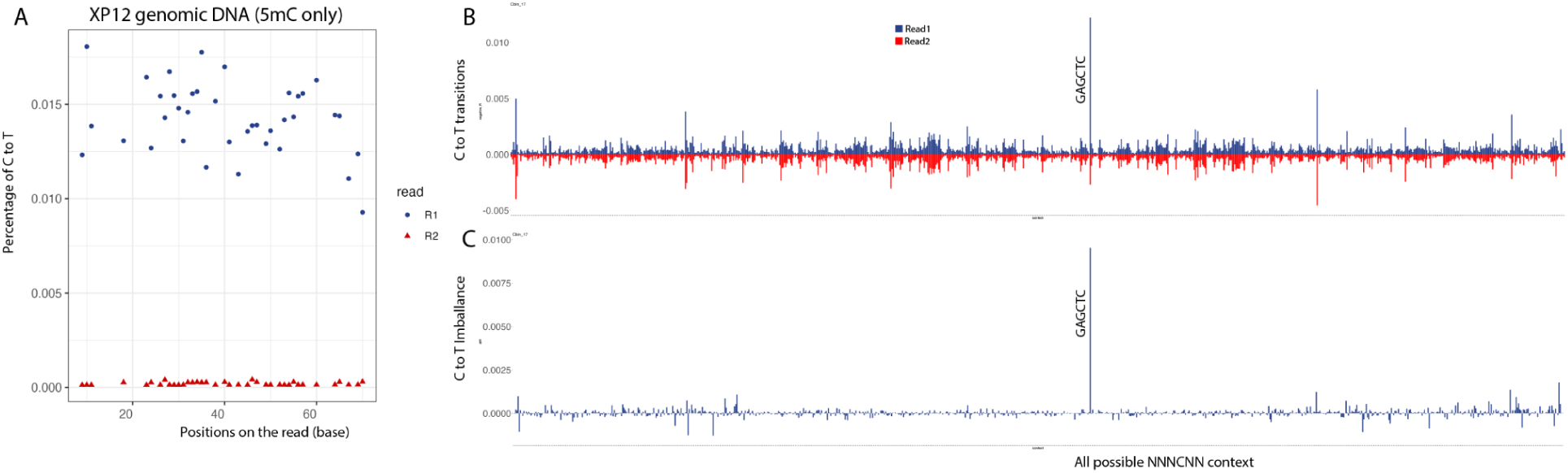
**A.** Heat-Alkaline deamination rate at ^m5^C genomic position (from the spike-in XP12 genomic DNA control). The C-to-T transition rate (in percentage) is calculated for each position on read 1 (R1, blue) and read 2 (R2, red). An average of 1.42 % and 0.019 % of C-to-T transition is obtained in read 1 and read 2 respectively. **B.** Absolute C-to-T transition rate in read 1 (blue) and 2 (red) in MAGs 17 for all NNNCNN contexts. **C.** Differential C-to-T transition rate between read 1 and 2 (imbalance value) in MAGs 17 for all NNNCNN contexts.

For each genome in the mock community, we applied mosdi-discovery (Marschall and Rahmann 2009) as described previously (Baum et al. 2021) to de-novo identify motifs predicted to be methylated. Using this approach, we identified all the methylation motifs that were previously detected by RIMS-seq and validated by bisulfite sequencing (**Supplementary Material**, **Supplementary Table 1**). Additionally, motifs were discovered with significantly fewer reads than required by RIMS-seq. For instance, the GAT<u>C</u> motif was identified with only 46,000 reads mapping to the *Bifidobacterium adolescentis* genome (**Supplementary Table 1**).

These results indicate that RIMS-seq2 can replace RIMS-seq for identifying methyltransferase specificities in mock microbiomes composed of a limited number of bacterial isolates at a fraction of the sequencing depth. However, it is important to note that the mock gut microbiome used in this study consists of an equimolar mixture of a limited number of bacterial isolates with available high-quality reference genomes. Consequently, this setup is not an accurate representation of native microbiomes.

### Proxi-RIMS-seq2 identifies contexts of elevated C-to-T transitions across phased-resolved microbiomes

Next, we performed RIMS-seq2 on two distinct native microbiomes: oral and vermicompost microbiomes, for which shotgun sequencing and proximity ligation (ProxiMeta) were performed to resolve assemblies into genomes (Proxi-RIMS-seq2). Additionally, we evaluated a reference fecal microbiome (ZymoBIOMICS TruMatrix™ Fecal Reference) that was collected from healthy adult donors and homogenized in one large batch. This reference composite microbiome was sequenced using PacBio and proximity ligation to obtain a high-quality, genome-resolved reference fecal microbiome (Portik et al. 2024). For all experiments, we aimed for an estimated 1% deamination rate at methylated sites (**Figure 1A, Material and Methods**). Proximity ligation libraries were produced using Phase Genomics’ ProxiMeta kits and the resulting libraries were sequenced on the Illumina NovaSeqX platform. Metagenomic assembly was performed using MEGAHIT and MAG deconvolution & analysis was performed using the ProxiMeta platform, with phage and plasmid genomes being reconstructed as described (Uritskiy et al. 2021). Sequencing of RIMS-seq2 libraries yielded approximately 200 million read pairs for the vermicompost, 160 million read pairs for the dental microbiome and 700 million read pairs for the TruMatrix dataset. RIMS-seq2 reads were mapped to the genome-resolved reference sequences using BWA-MEM (H. Li and Durbin 2009).

**Figure 1:**
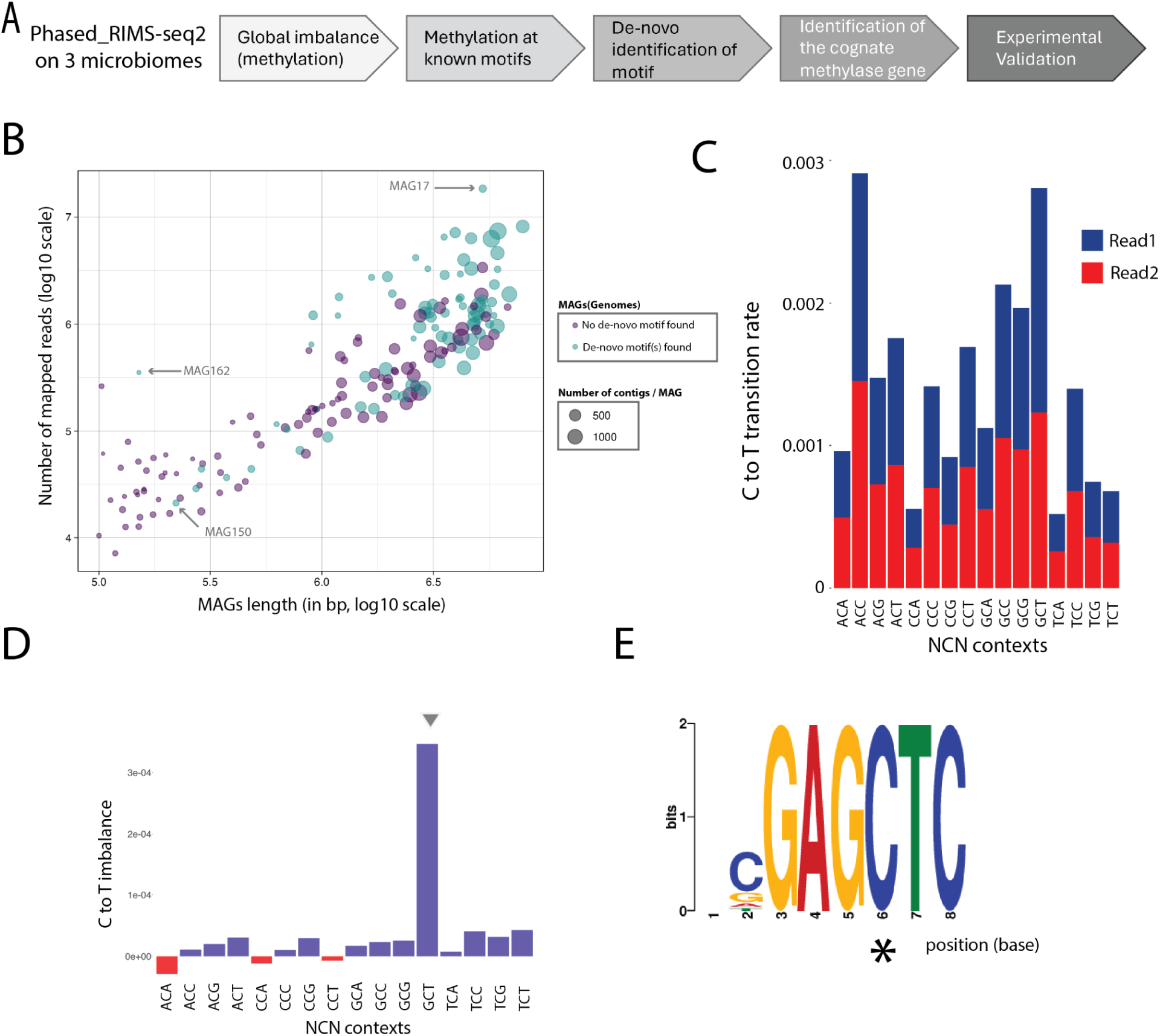
**A.** Overview of the Proxi-RIMS-seq2 analytic steps : [1] Imbalances in MAGs indicate methylation [2] imbalance at known motifs [3] imbalance is used to de-novo identify methylated motifs [4] linking genotype (MAGs) with phenotype (methylation) associates methyltransferases to their predicted recognition motifs [5] methyltransferase activities can be cloned and expressed in-vivo in strains lacking ^m5^C methylation for validation. **B.** Relationship between MAG length (in base pairs, x-axis) and read coverage (log10 scale, y-axis) within the genome-resolved vermicompost microbiome. Each data point represents a MAG, distinguished by the presence (green) or absence (purple) of predicted methylated motifs. Notably, MAG 17, belonging to the genus Patulibacter, exhibits the highest read coverage mapped to its consensus genome. **C.** Barplot representing the absolute C-to-T transition rate in read 1 (blue) and 2 (red) at all 16 N<u>C</u>N contexts(with N=A,T,C or G) in MAG 17. **D.** differential C-to-T transition rate between read 1 and 2 (imbalance value) at all 16 N<u>C</u>N contexts in MAG 17. **E.** PWM found most significantly associated with C-to-T imbalance in MAG 17 (* represent the methylated cytosine).

Using the vermicompost microbiome, we first assessed whether the imbalance between the C-to-T transition on read 1 compared to read 2 in RIMS-seq2 libraries could be observed despite the high heterogeneity of sequences typically found in population consensus genomes. Imbalance has been previously shown to represent damage rather than variants from the reference sequence (L. Chen et al. 2017) and in RIMS-seq, an imbalance of C-to-T represents a deamination of 5mC. Given that a typical microbiome is highly complex, we focused on a representative of the *Patulibacter* genus, whose assembly is 92% complete and has the highest read coverage in the RIMS-seq2 sequencing dataset (**Figure 1B**). Compared to mock microbiomes, we observed relatively higher C-to-T transition rates in all 3 base contexts (NCN with N=A,T,C or G) for both read1 and read2 (**Figure 1C**), presumably due to the heterogeneity captured in the population consensus genome. Despite this higher baseline variant rate, a prominent imbalance was observed in the GCT context (**Figure 1D**), indicating methylation in this context.

Next, we searched the *Patulibacter* genome for genes containing a ^m5^C methyltransferase domain using HMMER (Eddy 1998). A single hit containing a ^m5^C methyltransferase domain was found in the assembly. A homology search in REBASE (Roberts et al. 2023) identified the closest hits as M.Sgr13350I Type II methyltransferase with known activity at GAG<u>C</u>TC sites (Roberts et al. 2023). Knowing the expected motif content, we then sought to analyze the imbalance in all 6-base NNNCNN contexts. The result reveals significant imbalance only in the GAG<u>C</u>TC context at approximately 1% (**Supplementary Figure 1B and C**). This result is consistent with the expected deamination rate for RIMS-seq2 and the predicted methyltransferase specificity in *Patulibacter*. Finally, to evaluate whether such motifs could have been identified *de-novo* without prior knowledge of the predicted methylation motif, we used DiNAMO, a motif-searching algorithm that uses an exact discriminative method for discovering IUPAC motifs in DNA sequences (Saad et al., 2018). DiNAMO was supplied with all 15 bp sequences flanking a C-to-T transition events in read 1 (foreground) and read 2 (background) and identifies GAG<u>C</u>TC as the most significant motif associated with an excess of C-to-T transitions in read 1 (**Figure 1E**).

Taken together, these results indicate that C-to-T imbalance can be used to (1) assess whether a genome has ^m5^C methylation, (2) identify known methylated context(s), (3) *de-novo* identify methylated motif(s), and (4) predict the association between the identified motif(s) and their cognate methyltransferase genes in individual genomes recovered from metagenomes. While this has already been demonstrated for individual genomes, we have now established that a similar strategy can be employed directly from complex genome-resolved microbiomes, despite the high genotypic and phenotypic variability typically observed in population consensus genomes.

### Bacteria with similar methylation profiles tend to belong to the same order

Next, we examined the methylation profile of all three microbiome samples at 44 motifs known to be methylated by previously characterized methyltransferases from REBASE (see Materials and Methods). For this, we computed for each MAG the imbalance value in all these known motifs and subsequently clustered both the MAGs and motifs according to these imbalance profiles.

We observed that genomes within a cluster often belong to bacteria of the same order (**Figure 2** and **Supplementary Figure 2**). For instance, in the dental microbiome, methylation at AG<u>C</u>T motifs was predominantly found in the order Actinomycetales (**Figure 2B**). In the fecal microbiome, methylation at GGN<u>C</u>C and GGW<u>C</u>C motifs was commonly found in the order Lachnospirales (**Figure 2C**). While many methylation profiles are shared by related bacteria of the same order, other profiles were observed among evolutionarily divergent bacteria. For example, C<u>C</u>WGG and C<u>C</u>NGG contexts were methylated in a broad range of evolutionary diverse bacteria in the vermicompost (**Figure 2A).**

**Figure 2:**
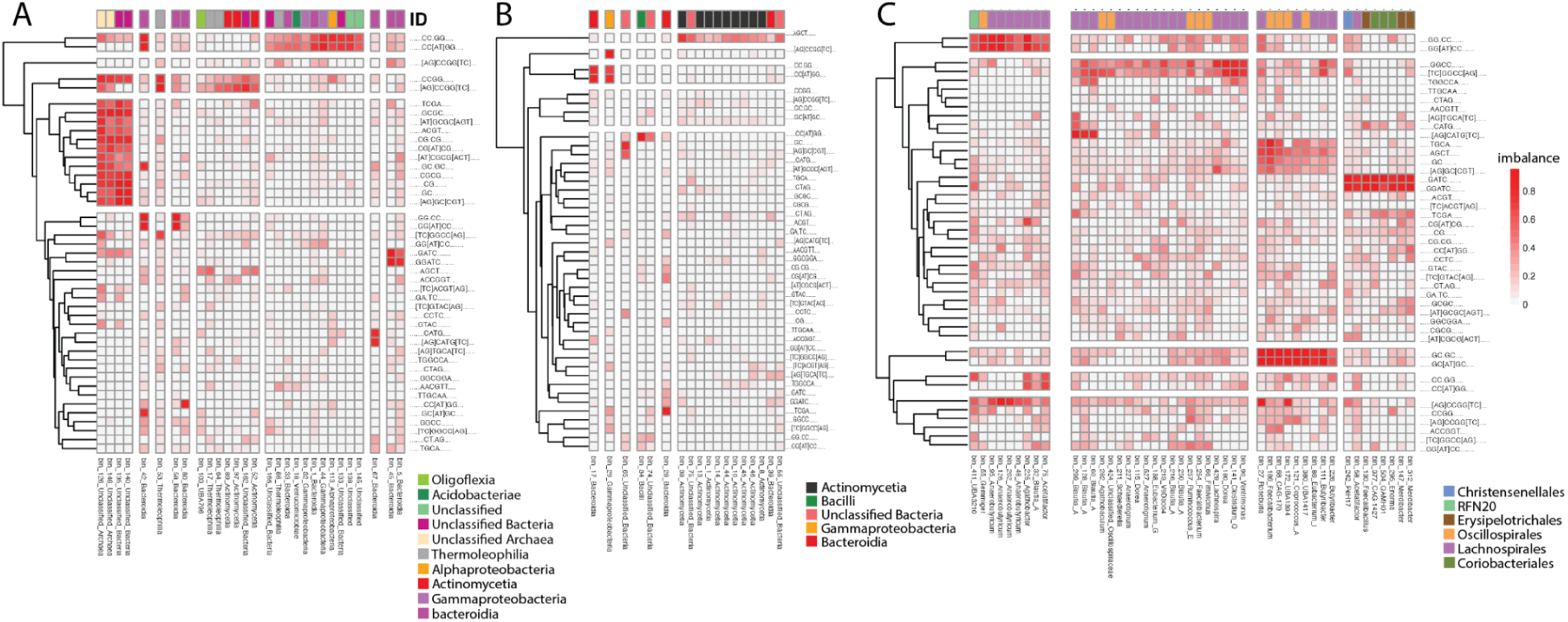
Imbalance profiles for selected clusters (full dataset in **Supplementary Figure 2**) in **A.** Vermicompost. **B.** Dental and **C.** TruMatrix microbiome. At an imbalance of 1% (dark red), all cytosine at the underlined position are predicted to be methylated. The x-axis corresponds to the annotated MAGs, y-axis corresponds to known recognition sites of ^m5^C methyltransferases (as cataloged in REBASE) and z-axis (gray to red) indicates imbalance values, ranging from 0 to 1%. MAGs are color-coded according to their taxonomic order. Both MAGs and motifs are clustered based on their imbalance profiles. Rows are clustered using the Manhattan distance, and columns are clustered using the Minkowski distance, as implemented in Pheatmap.

**Supplementary Figure 2:**
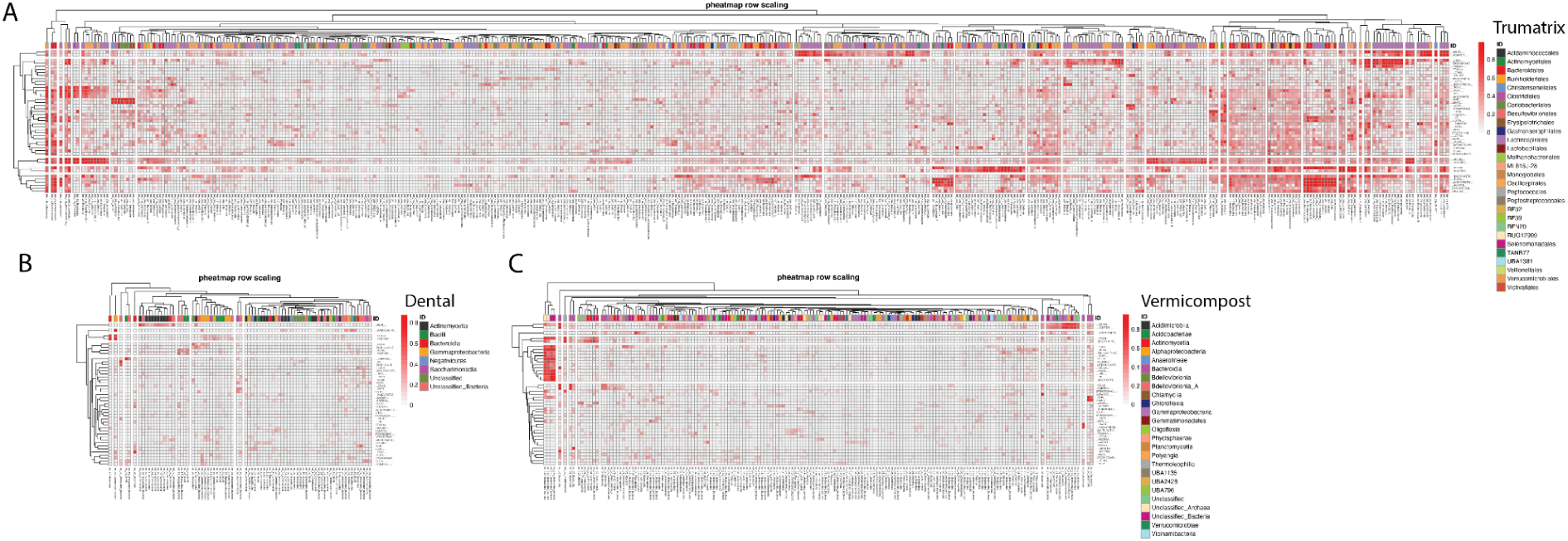
Imbalance profiles for all clusters. A. Vermicompost. B. Dental C. TruMatrix. At an imbalance of 1 (dark red), all cytosine at the underlined position are predicted to be methylated. The x-axis corresponds to the annotated MAGs, y-axis corresponds to known recognition sites of ^m5^C methyltransferases (as cataloged in REBASE) and z-axis (gray to red) indicates imbalance values, ranging from 0 to 1%. MAGs are color-coded according to their taxonomic order. Both MAGs and motifs are clustered based on their imbalance profiles. Rows are clustered using the Manhattan distance, and columns are clustered using the Minkowski distance, as implemented in Pheatmap.

### Proxi-RIMS-seq2 *de-novo* identifies ^m5^C methylated motifs and their cognate methyltransferases directly on native microbiomes

Next, for each MAGs, motifs associated with a C-to-T imbalance in the RIMS-seq2 paired-end libraries are identified using DiNAMO (**Material and Methods**). Accordingly, these motifs are predicted to be methylated in their respective MAGs. Most of the MAGs composed of assemblies of at least 1×10^6^ bp have at least one motif predicted to be methylated (**Figure 1B, Supplementary material**). Nonetheless, methylated motifs can be predicted in MAGs with as low as ∼20,000 reads (e.i, vermicompost, MAGs 150) or MAGs as small as 150,000 bp of total sequences. For example, with only 150,843 bp and 5.49% completeness, we were still able to predict two methylated non-overlapping motifs GC**<u>C</u>**GGC and GAG**<u>C</u>**TC in vermicompost MAGs 162.

A total of 75 motifs, 166 motifs and 1707 motifs were found in the 81 oral microbiome MAGs (0.92 motifs per MAGs), the 176 vermicompost MAGs (0.94 motifs per MAGs) and the 738 composite gut microbiome MAGs (2.3 motifs per MAGs) respectively (**Supplementary Material**). Motifs, methyltransferases and MAGs were deposited into Rebase (http://rebase.neb.com/rebase/rebase.html). We conducted several manual inspections of the link between Position Weight Matrix (PWM) and genotype data, often leading to successful matches even in complex cases involving multiple methyltransferase specificities. For example, we discovered five methylated motifs in MAG 4 with closest resemblance to *Algoriphagus terrigena* in the vermi-compost microbiome (**Supplementary Figure 2**). Six genes or gene fragments containing ^m5^C methyltransferase domains were identified on the *Algoriphagus terrigena* assemblies. We could assign with high confidence four of these genes to their cognate methylated motifs based on sequence identity with known ^m5^C methyltransferases for which the specificity has been experimentally verified (Roberts et al. 2023).

These results indicate that *de-novo* discovery of methylated motifs can be performed directly on native genome-resolved microbiomes, even when only a small fraction of the genomic sequences is available. Crucially, the binning of genomes allows for the linkage of genotypes to their epigenetic states, thereby enabling the association of methyltransferases to their predicted activities.

### Validation of newly identified methyltransferase specificities from Proxi-RIMS-seq2 data

The systematic association of novel methyltransferases with their sequence specificities at the microbiome level holds significant biotechnological potential, particularly in identifying enzymes with novel sequence specificities. To demonstrate the applicability of this strategy, we searched for novel motifs of interest that are predicted to be methylated.

One such newly identified methylation motifs is CAT<u>C</u>GATG which would corresponds to a novel 8-base pair recognition site. Methyltransferases recognizing such extended motifs are of significant biotechnological interest due to their potential association with restriction enzymes of similar specificity. This motif has been detected in two binned genomes from the vermicompost MAGs 10 and 51 corresponding to the genera *Oligoflexus* and *Anatilimnocola* respectively. Using HMMER (Eddy 1998) we searched for genes containing a ^m5^C methyltransferase domain and identified a single gene per binned genome. Pairwise comparison between *Oligoflexus* and *Anatilimnocola* methyltransferases shows 63.66% amino-acid identity across the entire protein and 68% identity across the Target Recognition Domain (TRD) which would suggest similar specificity. The closest methyltransferase homolog with known sequence specificities to those in *Oligoflexus* and *Anatilimnocola* is M.Bbr28III from *Bifidobacterium breve* with 48% and 52% identity respectively. M.Bbr28III has a known RT<u>C</u>GAY (with R = G or A and Y = C or T) motif specificity (partial of CAT<u>C</u>GATG) that has been confirmed by bisulfite sequencing (Roberts et al. 2023).

To experimentally validate the methylation specificities of these newly identified methyltransferases, we cloned and *in vivo* expressed them in an *E. coli* strain lacking endogenous ^m5^C methylation while conserving ^m6^A methylation at G<u>A</u>TC sites (NEB Express C2523 strain, dcm-; dam+). The *Oligoflexus* methyltransferase was successfully cloned and genomic DNA from the corresponding recombinant clone was sequenced using Tet-assisted PacBio SMRT-seq to assess methylation patterns (Clark et al. 2013). The result showed significant interpulse duration (IPD) ratio indicative of methylation at G<u>A</u>TC (^m6^A methylation from dam methyltransferase) and at CAT<u>C</u>GATG contexts (**Figure 3A-B**). Using the *de-novo* motif discovery tool DiNAMO, we were able to confirm both the ^m5^C modification CAT<u>C</u>GATG and ^m6^A G<u>A</u>TC motifs (**Figure 3C**) in the recombinant strain. These results demonstrate *Oligoflexus* ^m5^C methyltransferase activity at CAT<u>C</u>GATG as predicted by Proxi-RIMS-seq2.

**Figure 3:**
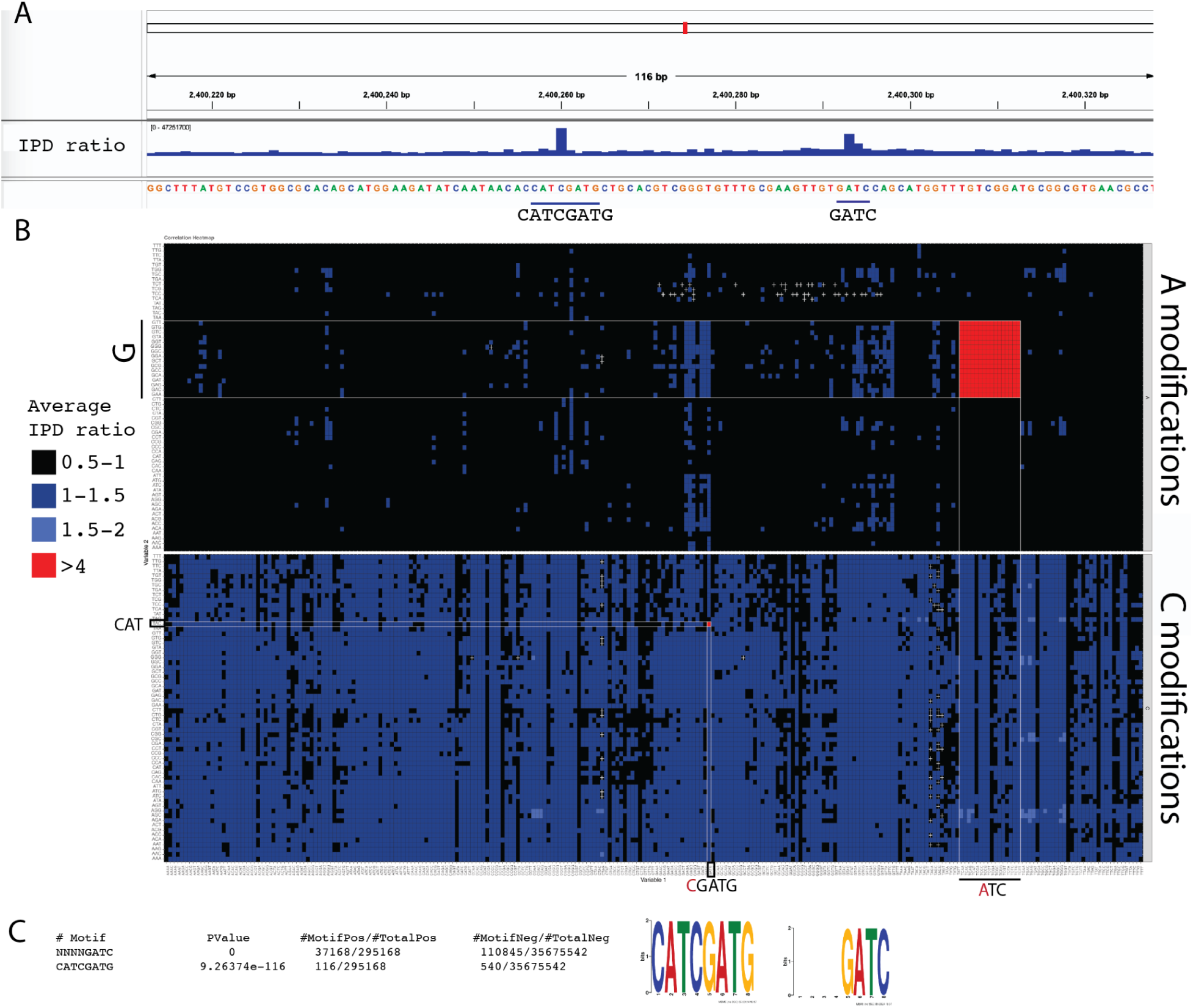
Validation of a new methyltransferase specificity using Tet-assisted PacBio sequencing. **A.** IPD ratios at a specific E.coli locus containing both CAT<u>C</u>GATG and G<u>A</u>TC motifs. **B.** average IPD ratio for all possible 8 mers for A modifications (^m6^A, upper panel) and C modifications (^m4^C or ^m5^C, lower panel). Y-axis corresponds to the 3 first nucleotides (oriented 3’ 5’), x-axis corresponds to the 4 last nucleotides (orientated 5’-3’) and z-axis (color) corresponds to the average IPD ratio. **C.** De-novo identification of motifs (output of DiNAMO and PWM). #MotifPos/#TotalPos :

### Proxi-RIMS-seq2 on viral DNA strengthened virus-host association

Proximity ligation provides linkage information to enable the association of viruses with their hosts. The physical interactions between phage DNA and the DNA of its microbial host are captured *in vivo* using proximity ligation, and the ProxiPhage algorithm (Uritskiy et al. 2021) is then used to reconstruct the vMAG and link it to its microbial host. We independently analyzed the predicted methylation profiles in both host and viral vermicompost MAGs and compared the identified motifs in each virus-host pair.

Out of 92 vMAG, 13 have at least one motif predicted to be methylated in both the MAGs and their corresponding vMAGs. Out of these 13 vMAGs, 10 have a good match with at least one motif from their predicted host (**Supplementary Figure 3**) and two have partial matches. These results indicate that the virus/hosts are sharing some of their methylation patterns and that the phasing of these interactions is reliable. In most of the cases, either the host or the virus have additional motifs possibly indicating a more complex stratification in the virus/host association than population consensus genomes can resolve.

**Supplementary Figure 3:**
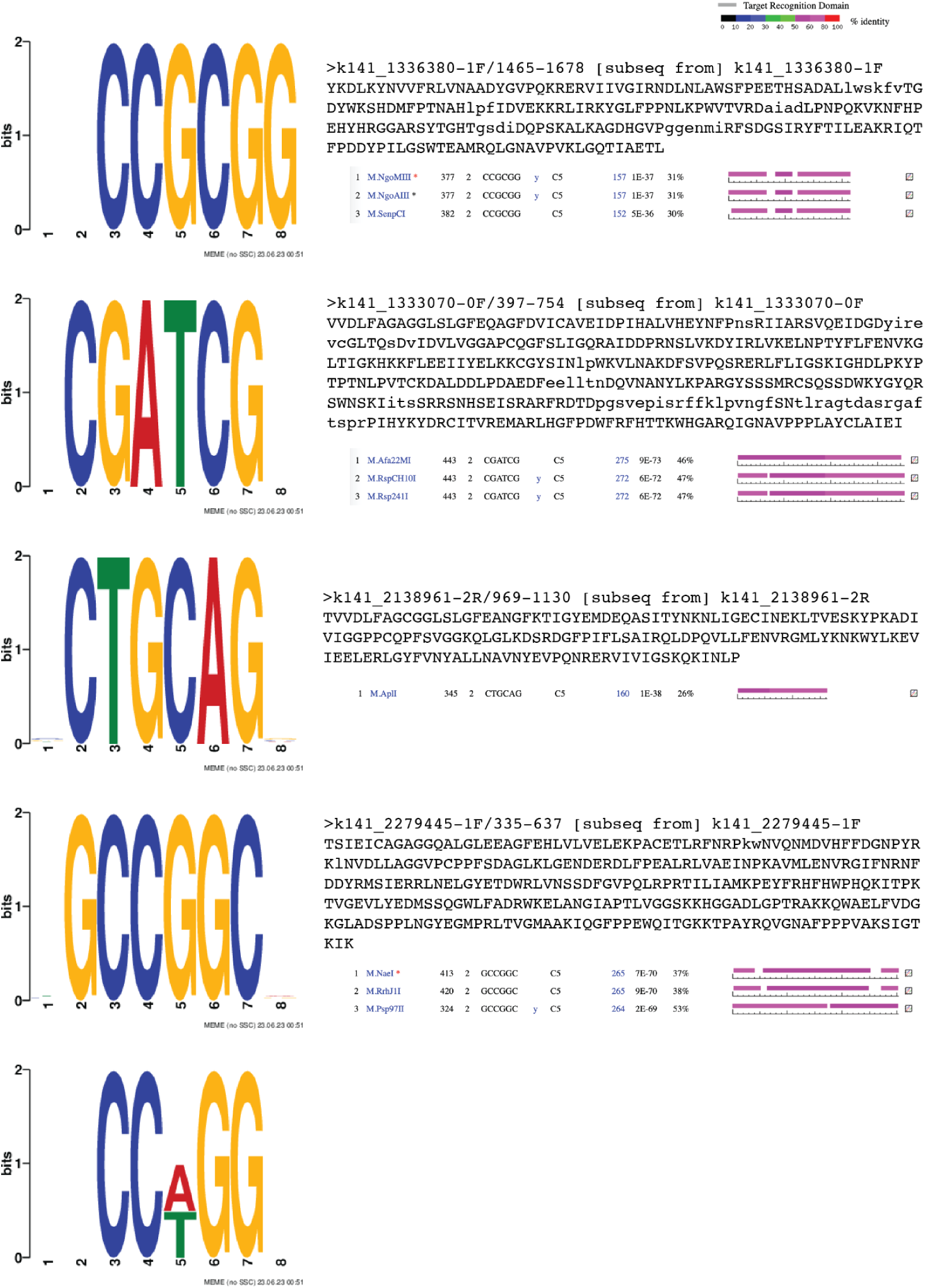
Example of multiple de-novo motifs (PWM) found in MAG 4 (closest hit: Algoriphagus terrigena) from the Vermicompost microbiome. The corresponding methyltransferase protein sequences are aligned to their predicted motif specificities. The correspondence between motifs/genes are predicted based on the closest hit in REBASE.

**Supplementary Figure 4:**
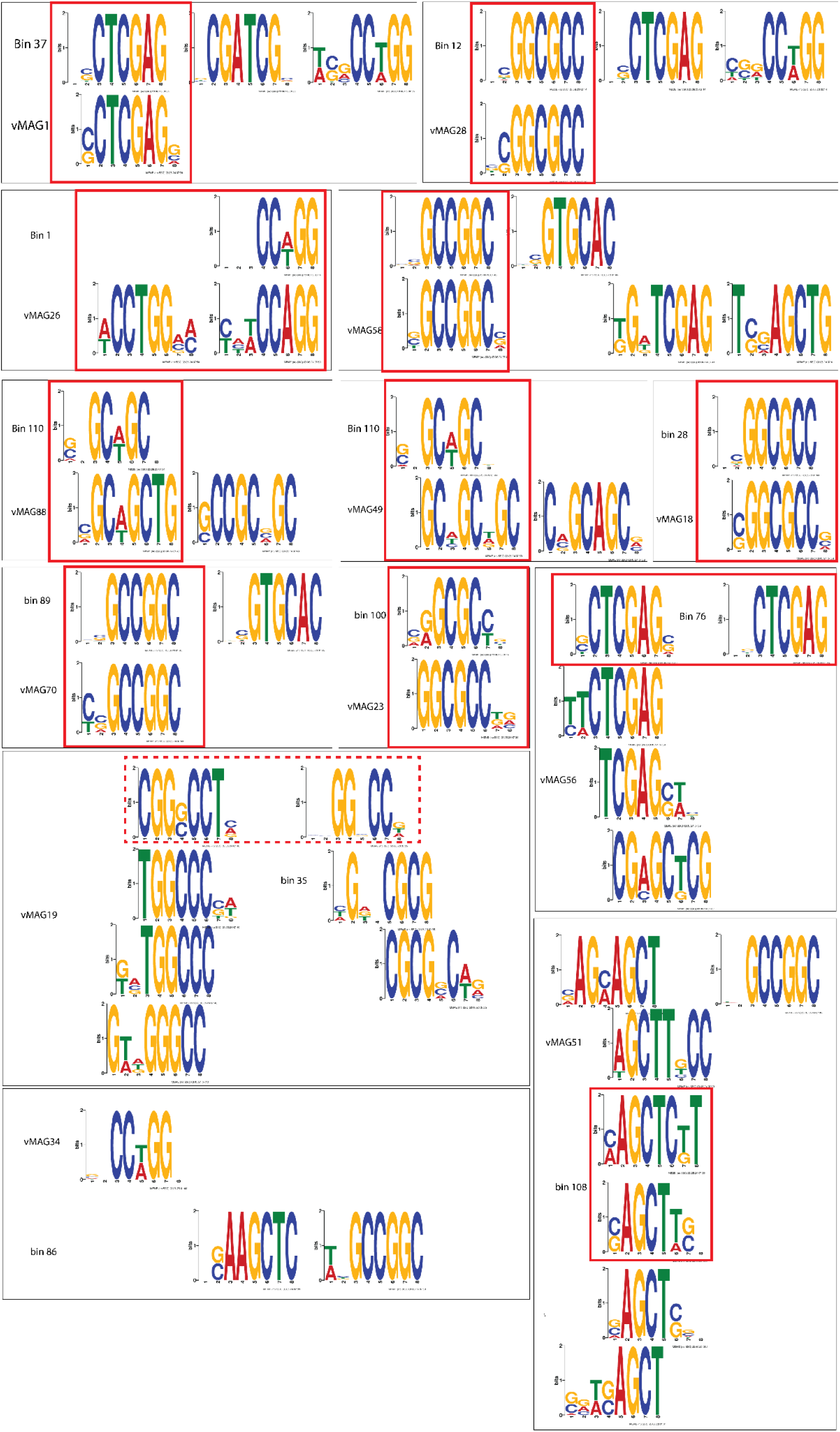
De-novo motif discovery in viruses (vMAGs) overlaid with motifs found in the corresponding host (MAGs). Red squares indicate good matches, while dotted red squares indicate reasonable matches.

## Conclusion

We developed Proxi-RIMS-seq2 to identify 5-methylcytosine (^m5^C) motifs directly in microbiomes and have demonstrated the ability to *de-novo* detect hundreds of such motifs across three distinct microbiomes. Two of these microbiomes were derived from unique samples, while the third, the TruMatrix microbiome, is a microbial reference material composed of stool samples collected from numerous healthy donors. Application of Proxi-RIMS-seq2 to the TruMatrix dataset revealed a higher number of methylation contexts per MAG compared with the other two microbiomes, suggesting a wide array of methyltransferase specificities. Given that TruMatrix is a meta-metagenome sourced from multiple donors, this observed diversity likely reflects the natural variability of methyltransferase genes across different individuals rather than an increased number or specificity of methyltransferase genes within a single stool.

Metagenome-Assembled Genomes represent population consensus genomes, and the degree of similarity in the epigenetic states within such populations remains unknown. Consequently, the observed methylation motifs likely originate from methyltransferases that are common and stable among a substantial portion of individual bacteria within these populations. Thus, the identified motifs in the studies may be biased towards stable epigenetic marks, such as the well-studied dcm methylation in *Escherichia coli* (Marinus and Morris 1973). Consistent with this statement, we often observed that methylated motif profiles are shared amongst related bacteria within the same order.

This study expands the potential applications of metagenomic data. For example, DNA methylation has been previously used as a complementary feature to enhance metagenomic binning (Beaulaurier et al. 2018). Similarly, Proxi-RIMS-seq2 can be employed to validate genome binning from proximity ligation data. Beyond improving binning accuracy, Proxi-RIMS-seq2 can be used to investigate the dynamics of methylation directly within microbiomes and to identify novel methyltransferase specificities, as demonstrated in this study with the identification of a new CAT^m5^CGATG recognition motif and its associated methyltransferases. This methyltransferase appears to function within a restriction-modification (RM) system, and we are currently characterizing the associated predicted restriction enzyme.

We have shown that it is possible to replace the conventional shotgun library with RIMS-seq while maintaining similar sequence accuracy and assembly statistics (Baum et al. 2021). Similarly, RIMS-seq2 could substitute for the conventional shotgun library that is required for the proximity ligation protocol. In this case we would achieve an efficient integration of methylation data with genome-resolved microbiome information, at minimal additional cost. Our study demonstrates that, despite decades spent searching for novel methyltransferase specificity for biotechnological applications, microbiomes still harbor hidden methyltransferase specificities yet to be discovered.

## Acknowledgments

We are grateful for the NGS core sequencing group at NEB for the sequencing of RIMS-seq libraries, Charles Elfe for providing a convenient download of REBASE, Tamas Vincze for incorporating the data to REBASE, Brian Anton for useful comments, Shuang-Yong Xu for advice and analysis of the 8 base-recognition restriction modification loci, Peter Weigele and Yian-Jiun Lee for providing Xp12 bacteriophage genomic DNA, Colleen Yancey for critical reading of the manuscript, and Alexey for advice on PacBio experiments. This work was supported in part by New England Biolabs, Inc., by NIH SBIR Grant R44AI172703 and a grant from the Bill & Melinda Gates Foundation to Phase Genomics.

## Data and code availability

All raw and processed sequencing data generated in this study will be submitted to the NCBI Gene Expression Omnibus upon publication (GEO; https://www.ncbi.nlm.nih.gov/geo/). Code for C-to-T imbalance calculation can be found in https://github.com/Ettwiller/RIMS-seq. MAGs, methyltransferase genes and motifs were deposited in REBASE (http://rebase.neb.com/rebase/rebase.html)

## Supplementary Materials

Supplementary Table 1 : QC and statistic on the mock microbiomes Supplementary material : Methylated motifs in microbiomes

## Materials and Methods

### Microbiome genomic DNA source

In this work, we performed RIMS-seq on four synthetic or native microbiome genomic DNA samples including: 1) mock gut microbiome genomic mix containing even mixture of fully sequenced and authenticated bacterial species observed in gut microbiome (ATCC, MSA-1006); 2) human oral microbiome (Phase Genomics); 3) human fecal microbiome (ZymoBIOMICS Fecal Reference, Zymo Cat #D6323) and 4) vermiculture microbiome (Phase Genomics).

### Genome-resolved microbiomes

A Hi-C library was created with the Phase Genomics ProxiMeta Hi-C v4.0 Kit using the manufacturer-provided protocol (Lieberman-Aiden et al. 2009). Briefly, intact cells from two samples were crosslinked using a formaldehyde solution, simultaneously digested using the Sau3AI and MluCI restriction enzymes, and proximity ligated with biotinylated nucleotides to create chimeric molecules composed of fragments from different regions of genomes that were physically proximal *in-vivo*. Proximity ligated DNA molecules were pulled down with streptavidin beads and processed into an Illumina-compatible sequencing library. Separately, using an aliquot of the original sample, DNA was extracted with a ZYMObiomics DNA miniprep kit (Zymo Research, Cat. #D4300) and a metagenomic shotgun library was prepared using ProxiMeta library preparation reagents. Sequencing was performed on an Illumina NovaSeq generating PE150 read pairs for both Hi-C and shotgun libraries. Hi-C and shotgun metagenomic sequencing files were uploaded to the Phase Genomics cloud-based bioinformatics portal for subsequent analysis.

Shotgun reads were filtered and trimmed for quality and normalized using fastp (S. Chen et al. 2018) and then assembled with MEGAHIT (D. Li et al. 2015)(D. Li et al. 2016) using default options. Hi-C reads were then aligned to the assembly following the Hi-C kit manufacturer’s recommendations (https://phasegenomics.github.io/2019/09/19/hic-alignment-and-qc.html). Briefly, reads were aligned using BWA-MEM (H. Li and Durbin 2010) with the −5SP options specified, and all other options default. SAMBLASTER (Faust and Hall 2014) was used to flag PCR duplicates, which were later excluded from analysis. Alignments were then filtered with Samtools (H. Li et al. 2009) using the -F 2304 filtering flag to remove non-primary and secondary alignments. Metagenome deconvolution was performed with ProxiMeta (Stewart et al. 2018) [10], resulting in the creation of putative genome and genome fragment clusters. Clusters were assessed for quality using CheckM (Parks et al. 2015) and assigned preliminary taxonomic classifications with Mash(Ondov et al. 2016).

### RIMS-seq library preparation and illumina sequencing

To prepare individual DNA libraries for RIMS-seq, we used gDNA input amounts ranging from 50ng - 80ng. The microbiome genomic DNAs were first sheared to 250bp with Covaris S2 Focused Ultrasonicator. One reaction of NEBNext Ultra II DNA Library Prep Kit for Illumina (NEB, #E7645) was used for each gDNA sample. Adaptor ligated libraries were purified with 1x volume of NEBNext Sample Purification Beads (NEB, #E7103) and eluted with 45 ul 0.1x TE buffer. NaOH was then added to the purified DNA libraries at 1M with a 30 min incubation time at 60°C followed by addition of an equimolar of acetic acid to neutralize the reaction. We included a USER (NEB, #M5505) treatment step to improve RIMS-seq library quality. Detailed protocol was published in our earlier work (Yan, Wang, and Ettwiller 2024).

Prepared libraries were amplified and indexed with NEBNext Multiple Oligos for Illumina (NEB, #E6446). Short-read sequencing was processed on an Illumina Nextseq instrument with paired end reads of 75bp.

Reads are mapped to the MAGs reference sequences using BWA-MEM (H. Li and Durbin 2009) and processed as described here (Baum et al. 2021).

### Imbalance in known contexts

A total of 44 IUPAC motifs, known to be recognition sites for at least 10 distinct ^m5^C methyltransferases cataloged in REBASE, were selected, and the positions of the methylated bases were recorded. The imbalance value was computed for all positions matching known motifs within a MAG. This value was calculated by subtracting the total number of C-to-T conversions in read 1 (R1) and G-to-A conversions in read 2 (R2) from the total number of C-to-T conversions in read 2 (R2) and G-to-A conversions in read 1 (R1), then normalizing to the total number of C-to-T conversions observed, using the following equation:

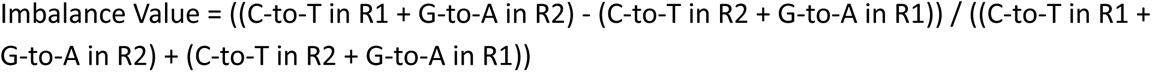

Instances of C-to-T or G-to-A conversions were identified using mpileup output. High-quality conversions were considered when the base quality score was ≥ 35 for C-to-T or G-to-A when compared to the reference assembly within a MAG. To avoid considering positions that contain true genetic variants, any position where the percentage of C-to-T or G-to-A conversions exceeded 5% for at least 5 reads was ignored. Programs and a detailed manual for the de-novo identification of motifs in Proxi-RIMS-seq2 are available on github (https://github.com/Ettwiller/RIMS-seq).

Clustering of the motifs and MAGs based on the imbalance values was done using the pheatmap package (version 1.0.12) with the following options : clustering_distance_rows = “manhattan”, clustering_distance_cols = “minkowski”.

### Proxi-RIMS-seq2 de-novo motif identification

Using the mpileup files, +/− 7 bp flanking genomic regions (15 bp total) for which a high quality (base quality score ≥ 35) C-to-T in R1 or G-to-A in R2 was found, were extracted for the foreground. Positions and +/− 7 bp flanking genomic regions (15 bp total) for which a high quality (base quality score ≥ 35) G-to-A in R1 or C-to-T in R2 was found, were extracted for the background. C-to-T or G-to-A in the first position of reads were ignored. If the percentage of C-to-T or G-to-A are above 5% for at least 5 reads at any given position, the position was ignored (to avoid considering positions containing true variants).

*De-novo* motif discovery in the mock microbiome has been performed using mosdi-discovery using the following parameters : ‘mosdi-discovery -v discovery -q x -i -T 1e-100 -M 8,2,0,4 8 occ-count’ using the foreground sequences with x being the output of the following command : ‘mosdi-utils count-qgrams -A ‘dna’’ using the background sequences. To identify additional motifs, the most significant motif found using mosdi-discovery is removed from the foreground and background sequences using the following parameter: ‘mosdi-utils cut-out-motif -M X’ and the motif discovery process is repeated until no significantly enriched motif can be found. A motif is found significantly enriched in the foreground sequence if P-value < 1e−100.

*De-novo* motif discovery in the microbiomes has been performed using DiNAMO. For each MAG, +/− 7 bp flanking genomic regions (15 bp total) for which a high quality (base quality score ≥ 35) C-to-T in R1 or G-to-A in R2 was found, were extracted for the foreground (file_foreground.fasta). Positions and +/− 7 bp flanking genomic regions (15 bp total) for which a high quality (base quality score ≥ 35) G-to-A in R1 or C-to-T in R2 was found, were extracted for the background (file_background.fasta). DiNAMO was run using the following parameters : dinamo -pf file_foreground.fasta -nf file_background.fasta -l 8 -t 1

### Cloning and expression of methyltransferase genes

Methyltransferase gene sequences were identified using the Hmmer (Eddy 1998) for the pfam domain PF00145.20 (C-5 cytosine-specific DNA methylase) (Mistry et al. 2021). Methyltransferase TRDs were identified using the conserved signature motifs identified by Posfai et al. (Pósfai et al. 1989)

Gene blocks for cloning methyltransferase genes were synthesized by Integrated DNA Technology (Coralville, IA). Molecular biology reagents including NEBuilder HiFi DNA assembly master mix, DNA size standards, competent cells, and NEBNext enzymatic methyl-seq reagents were provided by New England Biolabs (NEB). Genomic DNA isolation and plasmid purifications were performed using Monarch DNA kits (NEB).

The 1071bp methyltransferase gene was synthesized as 2 gene blocks (574bp and 608bp) containing overlaps designed for NEBuilder HiFi assembly. The expression plasmid pACYC184 (Chang, A.C. and Cohane, S.N. (1978) Construction and characterization of amplifiable multicopy DNA cloning vehicles derived from the P15A cryptic miniplasmid. *J. Bacteriol.,* **134,** 1141-1156) was amplified by inverse PCR. The DNA methyltransferase gene was cloned in the pACYC184 expression vector under control of a constitutive Tet promoter using NEBuilder HiFi DNA assembly master mix (#E2621) following manufacturer’s instructions and transformed into NEB Express (#C2523; g*fhuA2 [lon] ompT gal sulA11 R(mcr-73::miniTn10--*Tet^S^*)2 [dcm] R(zgb-210::Tn10--*Tet^S^*) endA1 Δ(mcrC-mrr)114::IS10*). Individual colonies were selected and grown overnight in LB broth supplemented with chloramphenicol (25μg/ml). Plasmid DNA and total DNA was isolated from overnight cultures using Monarch® plasmid mini-prep kit (NEB, #T1010) and Monarch® genomic DNA purification kit (NEB, #T3010) respectively. Plasmid DNAs were sequence verified by Oxford Nanopore Technology (ONT) sequencing using EPI2ME clone validation workflow (https://github.com/epi2me-labs/wf-clone-validation) and total DNA was prepared for sequencing on PacBio Sequel II (Pacific Bioscience, Menlo Park, CA).

### PacBio sequencing

*In vivo* modification activity and sequence specificity was analyzed by sequencing on the PacBio Sequel II. Prior to library preparation, input DNA was sheared to an average size of 5-10KB using gTubes (Covaris, Woburn, MA) and concentrated using 0.6V Ampure beads (Pacific Biosciences). Libraries were prepared using SMRT-bell Express Template Prep kit 2.0 (PacBio, #100-938-900) according to the manufacturer’s protocol. Barcoded libraries were TET-treated following an abbreviated protocol for NEBNext Enzymatic Methyl-Seq Conversion Module (NEB, #E7120) to enzymatically convert modified cytosines into 5-carboxylcytosine. Briefly, libraries were incubated for 1 hour at 37°C with TET2 in the presence of TET2 reaction buffer, an oxidative supplement, DTT, and an oxidation enhancer followed by addition of stop reagent and incubation at 37°C for 30 minutes. TET2 treated libraries were cleaned up using 1V Ampure beads (Pacific Biosciences).

TET-treated, barcoded libraries were prepared for sequencing on PacBio Sequel II using Sequel II sequencing kit 2.0 (PaBio, #101-820-200). Pacific Biosciences SMRTlink web portal was used to calculate sequencing conditions and perform post-sequencing analysis.

